# Explainable machine learning for health disparities: type 2 diabetes in the *All of Us* research program

**DOI:** 10.1101/2025.02.18.638789

**Authors:** Manoj S. Kambara, Onyinye Chukka, Kathryn J. Choi, Joseph Tsenum, Sonali Gupta, Nolan J. English, I. King Jordan, Leonardo Mariño-Ramírez

## Abstract

Type 2 diabetes (T2D) is a disease with high morbidity and mortality and a disproportionate impact on minority groups. Machine learning (ML) is increasingly used to characterize T2D risk factors; however, it has not been used to study T2D health disparities. Our objective was to use explainable ML methods to discover and characterize T2D health disparity risk factors. We applied SHapley Additive exPlanations (SHAP), a new class of explainable ML methods that provide interpretability to ML classifiers, to this end. ML classifiers were used to model T2D risk within and between self-identified race and ethnicity (SIRE) groups, and SHAP values were calculated to quantify the effect of T2D risk factors. We then stratified SHAP values by SIRE to quantify the effect of T2D risk factors on prevalence differences between groups. We found that ML classifiers (random forest, lightGBM, and XGBoost) accurately modeled T2D risk and recaptured the observed prevalence differences between SIRE groups. SHAP analysis showed the top seven most important T2D risk factors for all SIRE groups were the same, with the order of importance for features differing between groups. SHAP values stratified by SIRE showed that income, waist circumference, and education best explain the higher prevalence of T2D in the Black or African American group, compared to the White group, whereas income, education and triglycerides best explain the higher prevalence of T2D in the Hispanic or Latino group. This study demonstrates that explainable ML can be used to elucidate health disparity risk factors and quantify their group-specific effects.

**Author Summary:** While machine learning (ML) methods hold great promise for epidemiological studies, their practical utility is limited by interpretability. Increasingly complex ML models are great at predicting disease risk, but how they arrive at a given prediction is often obscured by model complexity. Explainable ML is an emerging discipline that seeks to render ML models more transparent by elucidating how and why input features contribute to output predictions. This study reports a novel application of explainable ML to epidemiology, focusing on type 2 diabetes (T2D) as a paradigm of health disparities. We found that ML classifiers were able to accurately model T2D disparities, for a large cohort of Black, Hispanic, and White Americans, and explainable ML revealed which risk factors contributed to the observed disparities and how. The results demonstrate that explainable ML can be a powerful tool for the discovery and characterization of health disparity risk factors.

## Introduction

Type 2 diabetes (T2D) is a highly prevalent disease with a larger impact on minority racial and ethnic groups in the US.(1-3) Black or African American and Hispanic or Latino individuals tend to be diagnosed with T2D at a higher rate compared to White individuals.(3-5) Studies have implicated a variety of physical, social, and environmental factors that are associated with T2D risk.(6-9) Traditional epidemiological studies focus on disease risk factors, i.e. variables that increase one’s risk of disease. We are interested in how these variables act as health disparity risk factors, i.e. risk factors that are associated with differences in disease prevalence between population groups.

Machine learning (ML) methodologies are increasingly being used for epidemiological studies of T2D.(10-14) Tree-based ML models in particular offer several advantages compared to regression methods that are more often used in epidemiology. Tree-based methods can utilize more complex model structures and incorporate more nonlinear covariate effects.(10, 15-17) Moreover, tree-based methods have been shown to have a higher sensitivity and specificity compared to regression models.(17) Tree-based models such as random forests (RF), extreme gradient boosted models (XGBoost or XGB), and light gradient boosting machines (lightGBM or LGB) are increasingly being applied in epidemiological studies.(10, 17)

Nevertheless, ML models are constrained by the “black-box” effect, which limits their interpretability. For disease classification models, it can be difficult to understand how individual features contribute to the final risk prediction. Explainable ML involves methodologies that address the “black-box” effect, thereby increasing the transparency and interpretability of ML models. SHAP (SHapley Additive exPlanations) values derived from cooperative game theory are a new class of methods that provide interpretability to ML models by quantifying the impact of each feature on the model’s predictions.(18) Algorithms calculating SHAP values leverage the hierarchical structure of tree-based models to track how predictions are influenced by transversing decision paths to compute exact SHAP values. In this way, SHAP values can illustrate whether a feature increases or decreases the likelihood of predicted disease risk. While methodologies of this kind are increasingly being applied in different disciplines, explainable ML methods have not yet been deployed to study racial and ethnic health disparities.

The goal of this study was to use ML methods and SHAP values to identify and quantify health disparity risk factors for T2D between Black and African American, Hispanic or Latino, and White participants from the *All of Us* Research Program cohort. *All of Us* is a large biomedical database of the US population, with participant data on demographics, genetic and environmental exposures, and health outcomes.(19-21) The first aim was to build and evaluate supervised ML models that accurately classify T2D risk and disparities among *All of Us* participants using a variety of risk factors. The second aim was to apply the methodology of SHAP values to quantify the effect of T2D risk factors on observed prevalence differences between racial and ethnic groups.

## Results

### Study participants and variables

We started with a cohort of n=287,012 participants that had EHR data. Of these participants we only included participants that could be classified as a T2D case or control based on Phecode 250.2 (n=219,728). Finally, we only included participants that had a SIRE of Black or African, Hispanic or Latino, or White (n=198,950). We used the full sample of n=198,950 for lightGBM and XGBoost modeling. Random forest modeling requires complete cases, giving us a smaller cohort of n=44,056. We visualized participant selection in **Supplementary Figure 1**.

We initially began with 26 putative T2D risk factors variables. We removed 2 variables (alanine aminotransferase, aspartate aminotransferase) based on high missingness, and 2 others (potassium, calcium) based on low effect sizes in LASSO regression (**Supplementary Figure 2**). **Table 1** shows the 22 variables included in ML modeling following selection. Variable selection is further described in the methods.

**Table 1.**
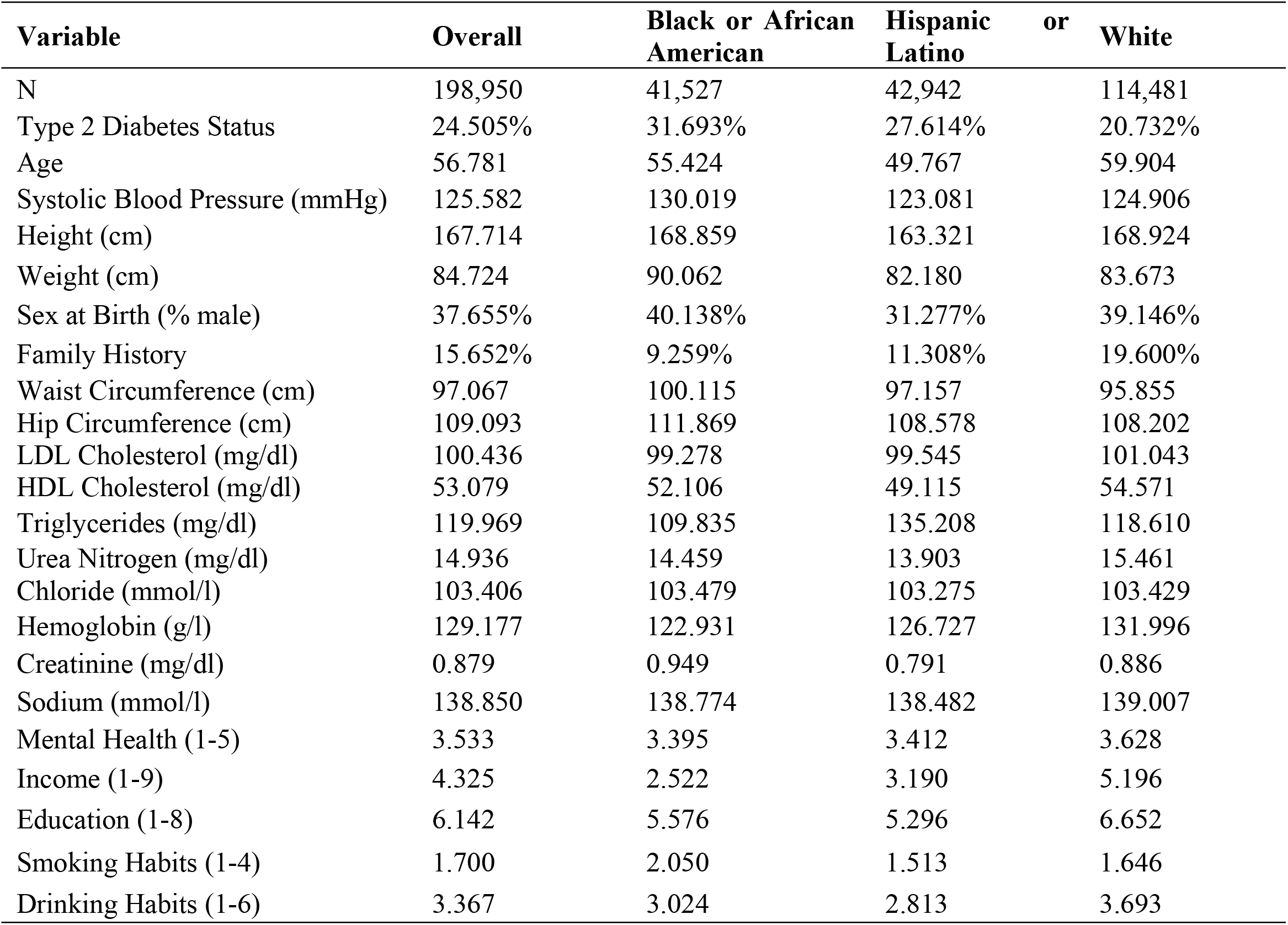
Participant cohort.

### All of Us reflects the US disparity in T2D

We found a clear disparity in T2D diagnoses between Black or African American, Hispanic or Latino, and White participants within the *All of Us* cohort, consistent with disparities of T2D in the US population. **Table 1** shows participant characteristics for the full cohort, including those with missing values. The overall T2D prevalence in the cohort was 24.505%. We further observe a disparity where White participants had the lowest prevalence of T2D (20.732%), followed by Hispanic or Latino participants (27.614%), and finally Black or African American participants with the highest prevalence (31.693%). We found the same trend in the complete cohort of participants with no missing data, as seen in **Supplementary Table 2** and **Figure 1**.

**Figure 1.**
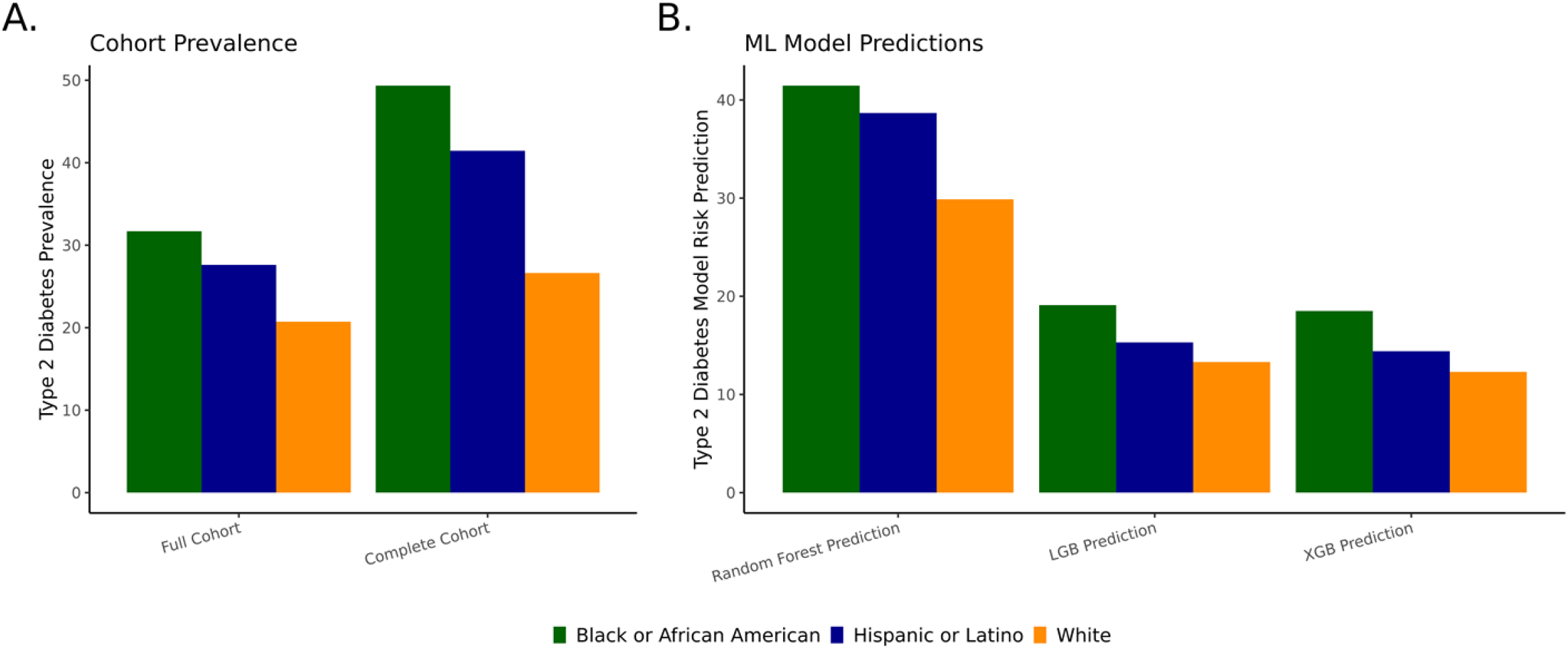
Cohort prevalence and ML model risk predictions. (A) Observed T2D prevalence for the *All of Us* cohort with complete cases (left) and cohort with missing data (right) for Black or African American (blue), and Hispanic or Latino (orange), and White (green). (B) ML model risk prediction for random forest (left), lightGBM (middle), and XGBoost (right) for Black or African American, Hispanic or Latino, and White participants. Calculated from sum of SHAP values for each SIRE group.

### Models classifying T2D are accurate

Three tree-based models, including random forest (RF), lightGBM (LGB), and XGBoost (XGB), were trained to classify T2D using 22 putative risk factors as model features, including blood biomarkers, demographic characteristics, lifestyle factors, physical measurements, and social determinants of health. All three ML models performed similarly well in classifying T2D cases and controls; all models exhibited high accuracy (*Accuracy*_*RF*_=0.797, *Accuracy*_*LGB*_=0.833, *Accuracy*_*XGB*_=0.831) and high AUCs (*AUC*_*RF*_=0.858, *AUC*_*LGB*_=0.870, *AUC*_*XGB*_=0.868). AUC-ROC curves for each model can be seen in **Supplementary Figure 3**. All models also performed similarly in regards to precision, recall, and F1 score. All metrics can be found in **Supplementary Table 3**.

### ML models recapture T2D disparities

For each model, we evaluated if the model was able to recapture the observed T2D disparities present in the *All of Us* cohort, namely showing the lowest predicted risk of T2D for White participants, followed by Hispanic or Latino participants, and the highest predicted risk for Black or African American participants. For each model, we calculated the SHAP value for each participant and took the mean of the SHAP values for Black and African American, Hispanic or Latino, White participants. We then evaluated the final risk prediction for each SIRE group. We found consistently that the ML models we developed predicted the lowest T2D risk for White participants, higher for Hispanic and Latino participants, the highest risk for Black or African American participants (**Figure 1**).

We chose to highlight the results from the random forest model because the SHAP values are calculated on a probability scale, lending to more clear interpretation. The random forest model had a high accuracy (0.797) and AUC (0.858). **Figure 2A, 2C, and 2E** shows the distribution of SHAP values for each variable given the mean value of the variable. We observed that the distribution of SHAP values tended to match the differences in variable values between each SIRE group. For example, mean age was lower for Hispanic and Latino participants (M_age,Hispanic or Latino_=54.981), compared to White (M_age,White_=62.456) and Black or African American (M_age,Black or African American_=58.932) participants. This was reflected by the higher density of negative SHAP values for Hispanic or Latino participants as seen in **Figure 2F. Figure 2B, 2D, and 2F** show waterfall plots for each SIRE group. Waterfall plots show the contribution of each feature to T2D risk prediction. As observed in **Figure 1**, the final risk prediction for each SIRE group matched the T2D health disparity observed in the cohort, with White participants showing the lowest predicted risk (0.299), followed by Hispanic or Latino participants (0.387), and the highest risk for Black or African American participants (0.415).

**Figure 2.**
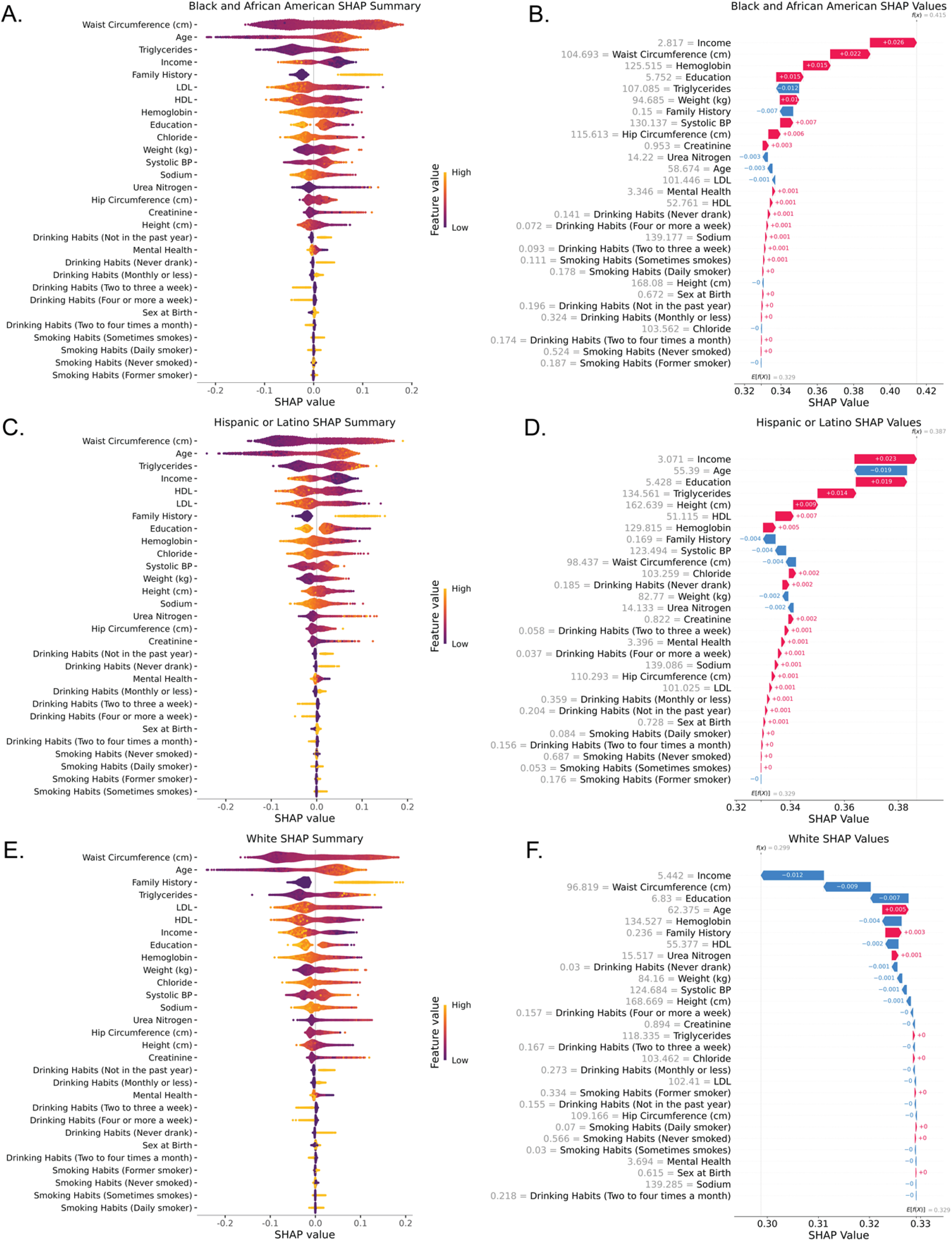
SHAP results for random forest model. (A, C, E) SHAP summary plots for each SIRE group. Shows how the SHAP value corresponds to feature value. Order of SHAP values is determined by feature importance, with the most important feature is on top. (B, D, F) Waterfall plots for each SIRE group. Feature SHAP values are the mean SHAP value for each feature in each SIRE group. Base risk predictions for all participants are shown as E[f(x)] and group-specific risk predictions are shown as f(x).

We observed the same trend in SHAP summary plots and waterfall plots for XGBoost and lightGBM models as seen in **Supplementary Figures 4 and 5** respectively. These models also captured a similar distribution of SHAP values and recaptured the health disparity. All models tended to treat features similarly as seen in correlations between SHAP values for each model in each SIRE group in **Supplementary Figure 6**, and through a comparison of feature importance between all models as seen in **Supplementary Figure 7**.

### Feature importance was similar for each SIRE group

As seen in **Figures 2A, 2C, and 2E** and **Supplementary Figure 7**, the top 7 most important features (based on mean absolute SHAP value) were common among all SIRE groups, with the order of importance differing between them. These features included waist circumference, age, family history, all cholesterol components (HDL, LDL, and triglycerides), education, income, and hemoglobin. Waist circumference and age were the most important features, respectively, for all SIRE groups. The other features differed in order of importance for each SIRE group.

### Group-specific feature importance explains difference in risk of T2D between SIRE groups

As seen in **Figures 2B, 2D, 2F** income was consistently the predictor with the largest magnitude for within-group risk of T2D. For white participants, income on average decreased T2D risk by - 0.012 (−1.2%), and increased risk for Black or African American participants by 0.026 (2.6%) and for Hispanic or Latino participants by 0.023 (2.3%). This was consistent with the difference in average income between groups, with White participants on average having a higher income comparatively to Black or African American and Hispanic or Latino participants. This indicates that income may be protective against T2D.

Education was also a predictor with a large magnitude for risk of T2D. For white participants, education slightly decreased T2D risk by -0.007 (−0.7%) and increased risk for Black or African American participants by 0.015 (1.5%) and for Hispanic or Latino participants by 0.019 (1.9%). This indicates that education may be another protective factor against T2D.

In addition to being the most important predictor for each SIRE group, we observed that waist circumference was another feature that greatly modified risk. For white participants, waist circumference slightly decreased T2D risk by -0.009 (−0.9%) and increased risk for Black or African American participants by 0.022 (2.2%) and decreased risk for Hispanic or Latino participants by -0.004 (−0.4%). Furthermore, the magnitude of SHAP values for waist circumference was consistently larger than the SHAP values for weight, indicating more importance in the location of fat on the body rather than simply weight.

Hemoglobin was also a predictor with a large magnitude for risk of T2D. For white participants, hemoglobin slightly decreased T2D risk by -0.004 (−0.4%) and increased risk for Black or African American participants by 0.015 (1.5%) and for Hispanic or Latino participants by 0.005 (0.5%). This indicates that elevate blood hemoglobin levels may be another protective factor against T2D.

## Discussion

ML methods are increasingly being applied in epidemiological studies. (10-14) SHAP values offer an avenue to uncover the traditional “black-box” of ML models and allow us to better utilize these models in the context of understanding disease. In this study we present a novel methodology of applying local SHAP explanations to the group level to identify and quantify health disparity risk factors for T2D. While the risk factors for T2D are well known, it is less known how these variables act as health disparity risk factors in contributing to T2D disparities.

We built multiple tree-based ML models to predict T2D risk using 22 putative risk factors as model features, including blood biomarkers, demographic characteristics, lifestyle factors, physical measurements, and social determinants of health. All models we developed showed a high accuracy and AUC in classifying T2D case-control status. Furthermore, the models we built effectively recaptured the disparity observed in *All of Us* and the US population.(3-5) Our models predicted the lowest risk for White participants, followed by Hispanic or Latino participants, and the highest risk for Black or African American participants.

Using these models, we calculated SHAP values for each individual and applied these individual-level explanations to the group level for each SIRE group. By doing this we were able to quantify the direction and magnitude of each feature’s contribution to T2D risk for each SIRE group. We found that socioeconomic factors, namely income and education, were among the strongest predictors of risk for each SIRE group. It is already well documented that these features play a role as health disparity risk factors in T2D.(6, 9, 32, 33) However, SHAP values reflect the directionality and magnitude of a feature’s contribution to overall risk, allowing us to quantify the observed health disparity risk factor. In our model, SHAP values helped us observe the relatively large contribution of income and education on T2D risk. SHAP values reflected how the differences in these variables between SIRE groups contributed to the overall within group risk for T2D. This framework shows how applying local SHAP explanations to the group level can be used as a means to better understand health disparity risk factors and understand health disparities using ML methods.

### Limitations

While we were able to strongly quantify some factors contributing to T2D health disparities using ML and SHAP values, there are limitations to consider. In regards to the cohort, the random forest model required participants with complete data, reducing the sample size. Furthermore, there was a clear unhealthy bias for participants from the Black or African American and Hispanic or Latino SIRE groups. When subsetting to complete cases, the T2D prevalence for these groups increased substantially. However, it is important to note that the disparity, i.e. the relative T2D prevalence between groups, was still comparable to the disparities observed in the US.(3-5) In regards to variable selection, while our list of variables was quite extensive, there is a possibility that we did not include a variable that is contributing to the T2D health disparity. In regards to the SHAP methodology, there are some limitations to consider. Firstly, SHAP values may not capture causal relationships, but rather correlations within the data.(34) Second, SHAP values for XGB and LGB models were on a log-odds scale, making the interpretability of those models more difficult. However, we showed that all models treated features the same. Finally, SHAP values assume that features within the model are independent of each other.(18) This means that the SHAP values may be missing an interaction between variables. This can be addressed in a future study by using SHAP interactions.(35)

### Generalizability

There are a few considerations to make regarding the generalizability of our results. Firstly, we make use of the *All of Us* cohort, which is comprised of volunteers. The *All of Us* cohort is older, more female, more educated, and does not match the racial and ethnic makeup of the US.(28-30) While we had a large, diverse sample of individuals within this study, the lack of representativeness may affect the overall external validity of the results.

## Conclusion

In conclusion, our study presents a novel methodology of applying explainable ML to characterize and quantify health disparity risk factors on the group level using SHAP values. Our study also highlights and reaffirms the large contribution of socioeconomic factors in health disparities for T2D. Further research can expand on the SHAP value methodology by factoring in interactions between features. Furthermore, as more data becomes available, this methodology could be applied to understand disparities in other groups and for other conditions.

## Methods

### Study Design

This study was comprised of two phases. The first phase involved building models that classify T2D using variables listed in **Supplementary Table 1**. To do this we used three tree-based ML models: random forest, extreme gradient boosting (XGBoost), and light gradient boosting machine (lightGBM). We built the random forest model in R using the randomForest R package.(22) We built the lightGBM model in R using the lightgbm R package.(23) We built the XGBoost model in Python using the xgboost Python library.(24) The ML models were trained to classify T2D status using 22 features listed in **Supplementary Table 1**.

The second phase of this study involved calculating SHAP values from the three models. To calculate SHAP values for random forest and lightGBM we used the treeshap package in R.(25) To calculate SHAP values for XGBoost we used TreeExplainer in Python.(18)

We used these SHAP values to understand disparities between Black or African American, Hispanic or Latino groups, and White participant SIRE groups. We stratified SHAP values by SIRE to identify and quantify health disparity risk factors. To do this we took the SHAP values for each participant in each SIRE group within the testing dataset and took the mean of the SHAP values for each feature. For example, we took SHAP values for White participants and then calculated the mean of each SHAP value for the entire group to get aggregate measures for each feature in the model. This approach is made possible by the fact that SHAP values are additive. After doing this we compared the relative SHAP value for each SIRE group and the differences in final risk prediction between each SIRE group.

### Setting

We utilized the *All of Us* Controlled Tier v7 dataset. This dataset can be accessed using the *All of Us* Researcher Workbench. More information regarding data collection for this dataset can be found here: https://support.researchallofus.org/hc/en-us/articles/14769699298324-v7-Curated-Data-Repository-CDR-Release-Notes-2022Q4R9-versions

### Participants

The cohort for this study consists of volunteers that have participated in the *All of Us* program. Volunteers can enroll for *All of Us* via JoinAllofUs.org or through a participating healthcare provider. Volunteers must reside in the US or in a US territory and be 18 years of age or older at the time of enrollment. Incarcerated individuals or those unable to provide consent cannot enroll in *All of Us*.

*All of Us* participant data was obtained from the *All of Us* Controlled Tier Dataset v7. We specifically obtained participant electronic health record (EHR), demographic, bloodwork, physical measurements, and survey data (**Supplementary Figure 1**). The period of enrollment for the Controller Tier Dataset v7 was from May 31^st^ 2017, to July 1^st^ 2022.

We initially selected participants that have contributed EHR data to *All of Us* (n=287,012). We then included participants that could be classified as a case (1) or control (0) for T2D based on Phecode 250.2 (n=219,728). Finally, we only included participants with a self-identified race and ethnicity (SIRE) of Black or African American, Hispanic or Latino, and White. The final cohort size was n=198,950. We included these three SIRE groups only because of the existing T2D health disparity between them and the fact that they are the largest groups in the *All of Us* cohort.(3-5)

Random forest modeling requires participants with complete data. From our final cohort, only n=44,056 had no missing data and were included in random forest modeling. LightGBM and XGBoost models can be built with incomplete or missing data, and such those models were built with the full cohort of n=198,950 participants.

### Variables

We defined T2D cases (1) and controls (0) based on Phecode 250.2.(26) This was done by mapping EHR data, specifically International Classification of Diseases codes (ICD-9-CM and ICD-10-CM) to Phecode 250.2.(26) If a participant had a Phecode between 249-250.99, we removed them due to having a condition similar to T2D.

We included 22 features to be included as predictors in classifying T2D. Clear definitions of all predictors can be found in **Supplementary Table 1**. We initially began with predictors from the American Diabetes Association Minute Test.(27) These are well-known and clinically relevant factors in T2D diagnosis. We further expanded this list to include other known factors in T2D diagnosis from other sources. This included adding waist and hip circumference, cholesterol (LDL, HDL, and triglycerides), income, education, smoking and drinking habits, and mental health.(6, 9) These features were chosen because they are documented in literature, were included within the *All of Us* dataset, and had low missingness within the *All of Us* cohort.

We further wanted to include other bloodwork factors with low missingness that were collected within the *All of Us* program. We initially included 7 bloodwork factors (urea nitrogen, chloride, hemoglobin, creatinine, sodium, potassium, and calcium). We conducted LASSO regression of these factors such that T2D status ∼ bloodwork factors. We selected the 5 bloodwork measurements with the highest absolute value of standardized beta coefficient: urea nitrogen, chloride, hemoglobin, creatinine, and sodium. Selection of bloodwork variables using LASSO regression is visualized in **Supplementary Figure 2** along with LASSO regression plots.

Many studies using ML to study T2D often include A1C or blood glucose as a predictor of T2D. (11-14) However, these factors are more indicative as outcomes of T2D, as they are used in T2D diagnosis. In order to understand differences in health disparity risk factors between SIRE groups, we wanted to focus on features that were predictors of T2D and thus, we excluded A1C and blood glucose. We also included social determinants of health known to be associated with T2D risk: education and income.

### Data sources/measurement

All outcomes and predictors can be found in **Supplementary Table 1**, along with sources highlighting variable selection.

### Bias

The *All of Us* cohort is comprised solely of volunteer participants, and thus, is not representative of the US population. The *All of Us* cohort has been shown to be older, more educated, more female, and does not match the racial and ethnic makeup of the US population.(28-30) This may affect the external validity of results from studies utilizing the *All of Us* cohort.

### Study size

The selection of participants can be seen in **Supplementary Figure 1**. The cohort for lightGBM and XGBoost was n=198,950. The cohort for random forest required complete cases, so the sample size was n=44,056.

### Quantitative variables

Quantitative variables were converted to ensure that there was consistency in units for each variable. For example, LDL cholesterol was not consistently measured in mg/dl. Data is recorded in mg/dl, g/l, and μg/l. To ensure consistency we converted all measurements to ensure that the measurements were in mg/dl. This was done for all quantitative features. The units for each variable can be seen in **Supplementary Table 1**. Quantitative variables were trimmed to only include values under the 99^th^ percentile. Variables above the 99^th^percentile were set to NA. This method ensured removal of abnormally large values.

### Statistical methods

Prior to model building we split the cohorts into training and testing datasets. We used an 80:20 training to testing split and stratified the split by T2D status to ensure the proportion of T2D cases was similar between both datasets. This ensured that proportion of cases and controls for T2D was similar between the training and testing datasets. This was done using the caret package in R.(31) These training and testing datasets were used in all model building.

All models were built such that the outcome was T2D status (control=0, case=1), and the features were all predictors included in **Supplementary Table 1**. Drinking and smoking habits were converted to dummy variables. Models did not include SIRE in the model. For each model, we evaluated the performance using the testing dataset. This included calculating accuracy, precision, recall, F1 score, and area under the curve (AUC) (**Supplementary Table 3**).

Following model building, we calculated SHAP values for each individual within the testing dataset for each model. Following this, we stratified participants by SIRE. Given the additive nature of SHAP values, we averaged SHAP values for each SIRE group and added them to get an average risk prediction as given by the model for each group. We evaluated the risk predictions to see if the models were recapturing the T2D health disparities in *All of Us*.

For the random forest model, we used only individuals with no missing data which included n=44,056. The training cohort included n=35,245, and testing included n=8,811. We built the random forest model in R using the randomForest package.(22) We calculated SHAP values for this model using the treeshap R package.(25) SHAP values for the random forest model were given in a probability scale.

For the lightGBM and XGBoost models, we used the full cohort including missing values which had a size of n=198,950. The training cohort included n=159,160 and the testing cohort included n=39,790. We built the lightGBM model in R using the lightgbm package(23) and calculated SHAP values using the treeshap R package.(25) We built the XGBoost model in Python using xgboost,(24) and calculated SHAP values using the TreeExplainer.(18) SHAP values for these models were given in a log-odds scale, thus we converted the final prediction to probability scale using the inverse logit function.

## Acknowledgements

The authors would like to thank the volunteers who participated in *All of Us*. Without them this study and the entire project would not be possible.

## Author Contributions

IKJ and LMR conceived of the study. LMR provided supervision and funding for the analysis in the *All of Us* Researcher Workbench. Most of the *All of Us* Researcher Workbench data analysis and the preparation of figures and tables were done by MK and OC. KC, SG, JT, NE contributed to the *All of Us* Researcher Workbench data analysis. MK and IKJ wrote and edited the manuscript. All authors read and approved of the final manuscript.

## Corresponding Author

Leonardo Mariño-Ramírez

11545 Rockville Pike Room C14, 2WF

Rockville, MD 20818

301-402-1366

## Data Access, Responsibility, and Analysis

All data and code for this analysis can be accessed via the *All of Us* Researcher Workbench. Individuals can apply for access at https://www.researchallofus.org/

## Abbreviations

AUC-ROC: Area Under the Receiver Operating Characteristic Curve LGB,
lightGBM: Light Gradient Boosting Machine Model
ML: Machine learning
RF: Random forest
SIRE: Self-Identified Race and Ethnicity
T2D: Type 2 Diabetes XGB,
XGBoost: Extreme Gradient Boosted Model

